# Transcriptomic Prediction of Breeding Values in Loblolly Pine

**DOI:** 10.1101/2023.03.21.533546

**Authors:** Adam R Festa, Ross Whetten

## Abstract

Phenotypic variation in forest trees can be partitioned into subsets controlled by genetic variation and by environmental factors, and heritability expressed as the proportion of total phenotypic variation attributed to genetic variation. Applied tree breeding programs can use matrices of relationships, based either on recorded pedigrees in structured breeding populations or on genotypes of molecular genetic markers, to model genetic covariation among related individuals and predict genetic values for individuals for whom no phenotypic measurements are available. This study tests the hypothesis that genetic covariation among individuals of similar genetic value will be reflected in shared patterns of gene expression. We collected gene expression data by high-throughput sequencing of RNA isolated from pooled seedlings from parents of known genetic value, and compared alternative approaches to data analysis to test this hypothesis. Selection of specific sets of transcripts increased the predictive power of models over that observed using all transcripts. Using information on presence of putative mutations in protein-coding sequences increased predictive accuracy for some traits but not for others. Known pedigree relationships are not required for this approach to modeling genetic variation, so it has potential to allow broader application of genetic covariance modeling to natural populations of forest trees.

## Introduction

Response to selection within breeding programs, commonly referred to as genetic gain, is defined as the change in population mean from one generation to the next due to artificial selection [1]. The amount of time required for one cycle of breeding, testing, and selection is called the generation interval and depends upon a species’ reproductive rate and the time a breeder waits to make future selections. For species with multi-year lifespans, breeders must wait until progeny are old enough to measure commercially-relevant traits in order to obtain accurate breeding value (BV) estimates with a high confidence level [2,3]. The appropriate age for measurement depends on the trait to be measured, but in some cases age-age correlations of phenotypes are high enough that traits measured at young ages are well-correlated with the same trait at an older age. Accelerating genetic gain by reducing generation intervals or establishing methods to reliably estimate individual BVs sooner could increase the value of any breeding program.

Traditional genetic evaluation in animal and plant breeding relies on phenotypic assessment of individuals and the genetic relationships between relatives. Given the recent increase of genomic resources within many breeding programs, utilizing sequence information to reduce the time estimating BVs has been explored and proven beneficial. One of the first examples of reducing time to estimating BVs through genomic information was genomic selection (GS), which produces dense marker coverage across a genome and takes advantage of associations between markers and phenotypes [4]. Since then, major annual crops such as maize (*Zea mays*), wheat (*Triticum aestivum*), rye (*Secale cereal*), and rice (*Oryza Sativa*) have shown improvement within their respective breeding programs through the application of genomic selection [5–8].

In addition, breeding programs for species that have longer generation intervals, such as dairy cattle and horses, have also shown that genomic selection is an efficient method for reducing the time of estimating BVs and thereby decreasing generation intervals [5–7]. Based on the efficiency of genomic selection and the constant reduction in sequencing costs, other ‘omic’ technologies have been explored. Omic methods are aimed at the detection of genes (genomics), mRNA (transcriptomics), proteins (proteomics), and metabolites (metabolomics) in a specific biological sample. These different types of data sets can be used separately or together to infer more about biological processes or as a means of accelerating the estimation of BVs and reducing generation intervals [8–17].

Forest tree breeding programs for species such as loblolly pine (*Pinus taeda*) are more time-consuming than for most crop species, because a single breeding and selection cycle takes roughly 15 years. Additionally, unlike most other breeding programs, tree breeding programs are in their infancy, typically no more than 3 or 4 generations deep with relatively weak pedigrees and connectedness among progeny test datasets [18]. Given this lengthy generation time and the relatively recent inception of tree breeding programs, the utilization of omic information in accelerating generation cycles and assisting in making future selections is an enticing opportunity.

At the moment, only a few experiments utilizing omic information for selection in forest trees have been conducted, all of which have focused on using either marker-assisted selection or GS [18–20]. Isik (2014) provides an excellent summary review of experiments that have been conducted in the field of forestry with regards to these types of tests. Due to the lengthy generation time of loblolly pine, a complete and robust experiment on utilizing genomic selection within the breeding program will take time to complete. A current experiment is underway to test the efficiency and accuracy of GS within the North Carolina State University (NCSU) Tree Improvement Program (TIP) breeding population using a base population of 2,300 cloned progeny from 53 crosses and is planned to be complete by 2032 [18].

The current selection method for advanced generations of loblolly pine breeding uses family mean and individual-tree phenotypes, where phenotypically superior individuals are selected from top-performing families [21]. Before a selected individual is used in a production population, progeny tests are carried out to confirm the genetic merit of the individual. These progeny tests are grown for four to six years and then measured for phenotypic traits such as volume, height, straightness, and fusiform rust disease incidence (caused by the fungus *Cronartium quercuum* f. sp. *fusiforme*). BVs of a tree for each trait are then estimated with linear mixed models where phenotype and pedigree data are utilized to help define the genetic covariance among a set of families from a mating design. The BVs determined from progeny tests are analyzed together with a pedigree of known relationships among individuals in the breeding population, using an algorithm to determine which selections should be used in the next breeding cycle to balance performance and amount of relatedness [22]. This requirement to progeny test all candidate selections increases the length of the breeding cycle. One attractive hypothesis is that “omic” selection methods may allow shortening of the breeding cycle, and increasing the rate of genetic gain per year, by allowing earlier selection of elite individuals and confirmation of the genetic value of those individuals.

It is reasonable to propose that genetic variation in family mean phenotypes can be partially accounted for by differences in gene structure and differences in gene regulation, given that heritable phenotypes of individuals are partly due to the underlying genetic architecture of a trait and genetic makeup within an individual. Published work in maize has reported accurate predictions of genetic differences among F1 hybrid progeny field performance based on pair-wise comparisons of gene expression levels among inbred parents of full-sib F1 hybrid families [23]. The authors identified a set of genes differentially expressed in at least one pair-wise comparison, then used gene expression data obtained from microarray experiments to create covariance matrices, either using the quantitative differences in expression or simple binary comparisons [23,24]. Utilizing a cross-validation approach with statistical models trained with a subset of the data, they confirmed that similarities in patterns of gene expression are correlated with similarities in field performance.

Loblolly pine does not have inbred lines or genetically-uniform hybrid F1 families; instead, the genetic entries used for deployment are either open-pollinated (OP) families of seedlings with a common seed parent, or control-pollinated (CP) families with both pollen and seed parents in common. An approach to transcriptomic prediction better suited to loblolly pine would be to use family-mean levels of gene expression values, because different loblolly pine families are known to have different genetic values for traits of interest. The most cost-effective method to estimate gene expression levels is by sequencing DNA copies of messenger RNA; this has been found to include less systematic bias when compared to microarray-based expression profiling due to the lack of background hybridization and has the additional benefit of identifying novel mRNA isoforms [25, 26]. Obtaining gene expression estimates from RNA-seq typically involves first fragmenting long RNA sequences and then converting the RNA fragments into short complementary DNA (cDNA) fragments. These fragments are then sequenced on a high-throughput instrument and aligned back to a transcriptome or genome to quantify the number of times a given gene or isoform has been detected. This initial quantification of gene expression estimates is systematically biased by many things, including the gene length, the batch that samples were sequenced in, and the lane or index used during the sequencing process [27]. To correct these systematic biases, transcript counts are typically normalized using various statistical approaches and then further used to address some hypothesis in a biological context. However, there is still no “gold standard” for normalizing the expression of RNA-seq reads accurately.

Genetic differences among families or individuals can be due either to differences in gene regulation patterns, or to differences in the structure of the encoded gene products [28]. Family-mean gene expression values obtained from RNA-seq may serve as a measure of the genetic covariance in gene regulation patterns. The DNA sequencing approach also provides information on DNA sequence variation such as single-nucleotide polymorphisms (SNPs) that can potentially reveal the genetic covariance among families with respect to gene structure. Utilizing these covariance structures for the prediction of parental BV’s, with or without feature selection of specific SNPs or specific transcripts, may provide predictive accuracy comparable to that obtained from family-based relationship models. Recent software tools allow for the incorporation of multiple covariance matrices, which may be used to assess these types of genomic information.

**The objectives** of this study were to 1) first assess the reproducibility of gene expression estimates obtained through sequencing replicate samples of seedlings from open-pollinated (OP), pollen-mix (PMX), or control-pollinated (CP) families. 2) We then aim to compare methods for utilizing family-mean estimates of gene expression to predict phenotypes. 3) Finally, we compare if predictive accuracy utilizing coding sequence SNP variation or gene expression level variation has independent value for modeling phenotypic variation or if they are redundant so that either data type (SNPs or gene expression) provides similar prediction accuracy.

## Materials and Methods

### Experimental design

#### Plant Material

Seedlings from 108 families (104 unique families) were planted in three separate batches, 46 families in batch one, 40 families in batch two, and 22 families in batch three over three different years. Families grown were produced from selections of the NC State University Cooperative Tree Improvement Program (NCSU-TIP) and the Western Gulf Forest Tree Improvement Program. Out of the 40 families planted in batch two, 15 families were selections made from the Western Gulf program. Across the first two batches, a total of four NCSU-TIP families were repeated. Seed used for families were obtained from a -20° C freezer stored at NCSU-TIP. The storage of seed is assumed to be the same for all families; however, the age of the seed differed. All families were stratified, sown, grown, and processed using the same conditions.

#### Seedling Growth

Stratification of seed was done by placing 150 seeds from each family in a plastic bag of water containing 1% hydrogen peroxide. They were stored at 4° C for 24 hours before draining and placed in moist bags at 4° C for 40 days. After stratification, seeds from each family were rinsed and planted into a subsection of a tray in a greenhouse located at NCSU in Raleigh, NC, USA. Trays were filled halfway with medium-coarse vermiculite as the soil medium. Prior to sowing seed, trays were lined with newspaper to prevent any leakage of vermiculite. After sowing a family on a subsection of the tray, a thin layer of vermiculite was spread on top of the tray to cover the seed after watering. The greenhouse was set to 24° C for all batches, and water was applied to seedlings three times a day.

#### Sample Collection

Families in the three separate batches were planted in independent years: batch one (September 2015), batch two (June 2016), and batch three (May 2018). Seedlings were allowed to grow for eight weeks after planting until a ‘rolling’ sample collection began. All samples within a given batch were harvested on a single day between 10 a.m. – 1 p.m., to minimize RNA expression differences due to time of day. Prior to collection, sample tubes for each family were labeled, where a single tube represents a single biological replicate of a given family. The number of biological replicates sampled depended on the number of seedlings germinated. If >90 seedlings germinated for a family, three biological replicates each containing 30 seedlings were collected. If <90 seedlings germinated, the biological replicates each consisted of 15 seedlings, and the number of replicates depended on the number of germinated seedlings. No other variation in the number of seedlings per biological replicate was allowed; for example, if 55 seedlings grew, only three replicates were taken to ensure comparable samples across replicates. Families that produced fewer than 15 seedlings were not collected. Over the first two batches, 77 out of the original 86 families had enough seedlings to collect samples. The first year contained 144 biological replicates from 46 families, and the second year contained 96 replicates from 31 families. All 22 families in batch three had enough samples to be collected and contained 66 biological replicates. A list of families sampled from each year and the numbers of biological replicates are represented in Supplementary Table 1. All biological replicates for an individual family were collected on the same day. Each replicate was collected using the following method:

- 30 or 15 seedlings from the tray subsection were gently removed from the planting mixture to retain roots and rinsed in water to remove vermiculite and debris
- Seedlings were quickly dried with a paper towel, placed in a mortar sitting on dry ice and ground with a pestle under liquid nitrogen
- A funnel prechilled on dry ice was then used to transfer ground seedlings to the pre-labeled prechilled tube sitting on dry ice.
- Tubes were stored on dry ice until transferred to a -80° C freezer.
- Between processing each family, the mortar, pestle, and funnel were wiped with 100% ethanol to prevent cross-contamination among families
- Before RNA extraction, all samples were ground using a coffee grinder, adding dry ice pellets to keep the tissue frozen. Each family was processed one at a time, and the coffee grinder was cleaned with 100% ethanol in between each family to minimize cross-contamination.

#### RNA Extraction and Sequencing

Once all samples were sufficiently ground, two independent RNA extractions were completed for each biological replicate, and the resulting RNA was pooled into one tube representing a single biological replicate. RNA extractions were done using a 96-well plate protocol modified from published methods for high-throughput DNA extraction [29].

For batch one and two, each pooled biological replicate was partitioned into three technical replicates and sequenced using multiplexing for batch one and two. Biological replicates were randomly assigned to one of 24 sequencing adapter index sequences, and technical replicates were randomly allocated to lanes. Each sequencing lane contained technical replicates with 24 different index sequences. The experimental design for each of the first two batches can be found in Supplementary Figure 1. Library creation, multiplexing, and Illumina HighSeq 2500 single-end sequencing were carried out by the Genomic Sciences Laboratory (GSL) at North Carolina State University in Raleigh, NC, USA. Batch three families were not partitioned into technical replicates; instead, biological replicates were sequenced directly. Library creation, multiplexing, and Illumina NovaSeq paired-end sequencing were carried out by Novogene at Davis, CA, USA.

Expression data received from the GSL and Novogene were processed using bbduk [30], clipping the first (left) 10 bases, filtering adapters, and quality trimming to a Phred20 score. Only the first read of each pair in batch three was used to make all batches as similar as possible. Trimmed reads were used to estimate gene expression levels on a biological replicate level using Sailfish [31] by aligning to a loblolly pine transcriptome produced at Indiana University by Don Gilbert [available at http://arthropods.eugenes.org/EvidentialGene/plants/pine/] (henceforth referred to as the 78k transcriptome because it contains about 78,000 putative transcript contigs).

#### Evaluation of expression similarity

Reproducibility of RNA-seq read counts per transcript across biological replicates and family mean values used spearman-rank correlation analyses. The Spearman correlation coefficient (*r*_*s*_) is defined as the Pearson correlation coefficient between a set of ranked variables. For *n* number of transcripts, the counts within each individual *X*_*i*_ and *Y*_*i*_ are converted to rank, *rgX*_*i*_ and *rgY*_*i*_ respectively. The Spearman correlation coefficient between the two individuals is then estimated as:

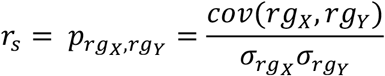

Where *p* is the Pearson correlation coefficient applied to the ranked variables, *cov*(*rg*_*X*_,*rg*_*Y*_) is the covariance of ranked variables, 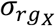 *and* 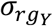 are standard deviations of the ranked variables. Unlike the Pearson correlation, rank-correlation is not as prone to being impacted by outliers, which reduces the impact of having multiple low or highly expressed transcripts on the resulting correlation coefficient. All correlation estimates were constructed using log-base-2 transformed counts.

#### SNP identification

Candidate SNPs were identified by aligning filtered and trimmed fastq files of all replicates across the three batches to the 78K loblolly pine transcriptome one at a time using bowtie2, producing individual SAM output files [32]. Samtools was used to convert SAM files to BAM files and sort them [33]. Picard was used to merge BAM files of biological replicates into a single BAM file representing one family replicate [34]. Bamaddrg was used to add read groups to files [35]. SNP calling was done using freebayes with settings to ignore indels, multi-nucleotide polymorphisms, complex events, and assuming diploidy of the samples [36]. SNPs reported in the VCF output from freebayes were further filtered using VCFtools to retain candidate SNPs with QUAL values > 30, minor allele frequency (MAF) > 0.05, and no missing data. The VCF file filtered for sites was processed with VCFtools to recode SNP genotypes into gene content 0,1,2 [37].

To evaluate the initial relatedness of families, the gene content matrix was converted to a standard genomic relationship matrix *G* using the VanRaden method [36].

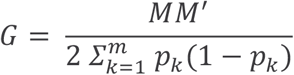

Where **M** was the minor allele frequency (MAF) adjusted genotype matrix with elements (0−2*p*_*j*_), (1−2*p*_*j*_), and (2−2*p*_*j*_) representing genotypes AA, AB, and BB, respectively; *p*_*j*_ was the MAF of the *jth* SNP. The construction of the *G* matrix was done using the AGHmatrix package in R, and a heat map of the resulting relationships was plotted [39,40].

### Prediction of Parental Breeding Values

Family empirical breeding values for volume were obtained from the NCSU TIP internal database. Breeding values for these parents were calculated from a wide range of testing sites and different trials. All prediction strategies used features from either SNPs or transcripts on the family-mean level and were conducted in R using the packages *OmicKriging* and *glmnet* [41,42].

The OmicKriging model implements a statistical method for optimal or best unbiased linear prediction by using a composite similarity matrix to make predictions based on the known relationships among other individuals [41]. The prediction of a test individual in OmicKriging is computed as the weighted average of the phenotype of individuals in the training set.

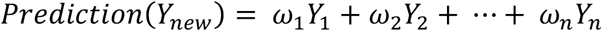

Where weights *ω*_*i*_ are defined as a function of all *n*(*n* + 1)/2 pairs of similarity measures, and Y corresponds to the individual phenotype. The weights prescribed by OmicKriging are defined as

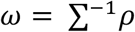

where ***ρ*** is a similarity vector between any test individual and the remaining training individuals and **Σ** is a similarity matrix of individuals within the training set. A single composite similarity matrix was used for prediction so that

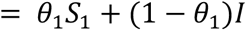

where the weights *θ’s* are supplied to each similarity matrix (*S*_*s*_) and **I** is the identity matrix used to represent an environmental component independent across individuals. All predictions with OmicKriging were conducted using a Pearson correlation matrix with a weighted value of 1.

In most genomic resources, highly correlated features and the need for feature reduction exist when n << p (where n is the number of samples and p is the number of features). To contrast OmicKriging with a model that intrinsically applies feature selection, we evaluated elastic net (EN) with the R package *glmnet* [42]. EN is referred to as a compromise between ridge and LASSO regression penalties. Ridge regression has the drawback of not performing feature selection and shrinks coefficients of correlated variables toward one another. At the same time, LASSO can only choose at most *n* variables and typically selects the strongest of highly correlated variables. Given that we expect transcript and SNP data to have a significant amount of highly correlated features and that most features are unrelated to phenotype, utilizing EN allows us to enable feature selection while not limiting ourselves to only *n* variables.

The glmnet model with EN contains two tuning parameters, alpha (*α)* and lambda (*λ)*, used to tune the penalty term

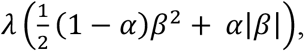

of the equation [42]:

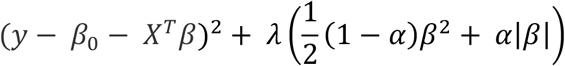

The parameter *α* controls the type of shrinkage, and parameter *λ* controls the amount of shrinkage. EN prediction was performed with the *caret* package in R [43]. In order to choose the tuning parameters, the training set was bootstrapped 25 times using a grid search with an alpha sequence of 0.1 to 0.9 by 0.1 and a lambda of 1, 0.1, 0.01, and 0.001 [43]. The best-tuned model on the training set, evaluated by the root mean squared error (RMSE), was used for prediction on the test set.

#### Creation of test and training sets

To assess the ability to predict family breeding values of seedlings grown across multiple batches, batch one was used as the training population to predict batch two and three as the test population. To evaluate if combining batches impacted prediction, individual predictions of batch two and three were conducted using batch one plus the other batch as training to predict the families in the left-out batch. In all cases, families that overlapped between batch one and two were used in the training set and not in the prediction set.

#### Prediction using transcripts

To make predictions on the family level, the mean of read counts across biological replicates corresponding to a single family were taken. The family-mean read counts were then added to one (to eliminate zero values) and log-transformed. To remove genes with low expression levels, we compared removing transcripts that had less than a mean of two, or a mean of three, family counts on the log2 scale. The resulting set of transcripts which passed this threshold were used as a baseline prediction in OmicKriging and glmnet (this set of predictions is referred to as ALL in result tables). A one-way ANOVA was performed to compare the batch effect on family-mean log2 transformed counts to account for batch effects. Individual families that were present in both batch one and two were removed prior to conducting ANOVA. The resulting p-values were adjusted using the lfdr function from the R package *qvalue* [44], and any feature with P < 0.05 was removed. Prediction with this set is referred to as setA in the results. To test if eliminating correlated features improved prediction, a correlation matrix of the batch effect filtered set of transcripts was evaluated with the findCorrelation function in the R package *caret*. Transcripts that explained more than 25% of the variation in another transcript were removed, and the resulting set was then used for prediction with *OmicKriging* and *glmnet*. In the results, prediction with removal of correlated features is referred to as setB. The Spearman correlation of transcripts in each training set against the phenotypes was performed to assess if features could be subset further. Transcripts that had an absolute correlation greater than 0.05 with phenotype were then included for prediction; these predictions are referred to as SetC.

#### Prediction using SNPs

All SNPs were used as the input to *glmnet* and *OmicKriging* as a baseline; these predictions are referred to as ALL in the results tables. To remove any SNPs affected by batch, a one-way ANOVA was performed to compare the effect of batch against the 0,1,2 family SNP matrix. The resulting p-values were adjusted using the lfdr function of the *qvalue* package, and any feature with q < 0.05 was removed. The relative variance was identified for each SNP by dividing the variance of SNP genotypes by the mean genotype for each locus. All SNP loci with values greater than 1 using this ratio were subset and used for prediction, referred to as setA. Correlated variables explaining 25% or more variation were removed and used to predict the test set; these results are referred to as setB. Additional filtering was explored by generating anovaScores using training data and phenotypes with the R package *caret*. Any value less than 0.05 was used for prediction; these results are referred to as setC.

#### Measure of accuracy

Predicted and observed breeding values were evaluated using simple linear regression and RMSE estimates between the two values to quantify prediction accuracy for a given model. A paired-samples t-test was performed comparing the absolute residual errors of prediction minus true breeding values between two models to see if a given model predicted families more accurately than another. The mean of prediction values within a given model was taken when evaluating the combination of transcripts and SNPs.

## Results

### Illumina Sequencing and Sequence Data Analysis

The samples from the first two batches yielded 3,639,060,450 126-nt reads, and samples from the third batch produced 1,444,563,627 150-nt reads. After trimming sequence adapters and low-quality regions, 2,605,756,024 reads (71.6 %) remained in batch one/two and 1,237,931,198 reads (85.7 %) in batch three. The average length of trimmed reads was 112 base pairs within batch one and two, and the average base quality was 37.1. The average length of trimmed reads for batch three was 140 base pairs, and the average base quality was 36.1.

Across all three batches, total read counts per biological replicate prior to applying any transformation ranged from 10.56 million to 39.24 million, with a mean of 20.76 million reads. For the 78,213 transcript contigs in the reference transcriptome assembly, read counts summed across all 255 biological replicates ranged from a minimum of 0 reads to a maximum of 294.96 million, with a mean of 67,688 reads. Out of the total 78,213 putative transcripts, a total of 40,942 (52.3%) transcripts passed the threshold of having a mean of at least two counts and 34,942 (44.7%) transcripts passed the threshold of having a mean of at least three on the log2 scale across all families. The log2 mean count across all putative transcripts within each of the 78 families ranged from 5.646 to 7.583 and standard deviations ranged from 2.408 to 2.759.

Approximately 3.92 million putative SNPs were initially identified from the merged bam files using freebayes. Filtering the candidate SNPs for those with a minor allele frequency no less than 0.05, a minimum Phred-scaled quality score of 30, and no missing data resulted in 214,449 candidates.

Out of the 306 biological replicates, 51 biological replicates were removed due to the following reasons: 15 biological replicates showed incorrect allele sharing relationships in a SNP relationship matrix, three biological replicates corresponding to a single family were missing an accurate phenotype, and 33 biological replicates represented selections from the WG TIP which were sequenced to be used in a separate study. These 51 biological replicates were removed, leaving 255 biological replicates comprised solely of NCSU TIP Coastal selections for analysis. Due to consolidation of some pollen mix and open-pollinated families, as well as the number of families that had enough germinated seedlings to prepare sequencing libraries, 78 families were used in prediction: 40 in batch one, 11 in batch two, of which five are present in both batch one and two, and 22 in batch three.

### Relationships of individuals using transcript or SNP data

To address the consistency in gene expression estimates among open-pollinated or control-pollinated families, correlations among biological replicates within each respective family were evaluated, ranging from 0.87 to 0.97 and 0.95 to 0.97 respectively (Figure 1). The similarity of gene expression estimates across batches grown in different years ranged from 0.89 to 0.94 (Figure 1). Correlation of biological replicates across all families ranged from 0.80 to 0.98 (Figure 1). The 3003 comparisons of family pairwise correlations showed overall gene expression patterns to be highly similar across families, with pairwise Spearman correlations ranging from 0.85 to 0.98 (Figure 1). Unlike transcripts, relationships among families using the 214,449 SNP genomic relationship matrix showed minimal relationships among most families, as expected (Figure 2).

**Figure 1.**
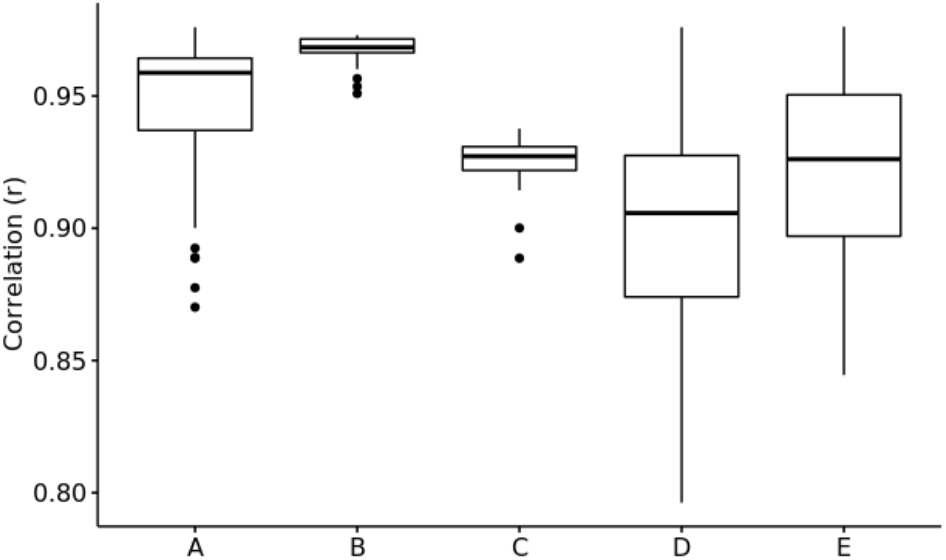
Spearman correlation (r) of gene expression estimates log2-transformed. (A) Open-pollinated biological replicates within family (N=329); (B) Control-pollinated biological replicates within family (N=23); (C) Biological replicates across batches (N=40); (D) Biological replicates within family (N=32385); and (E) Family-mean gene expression counts (N=3003).

**Figure 2.**
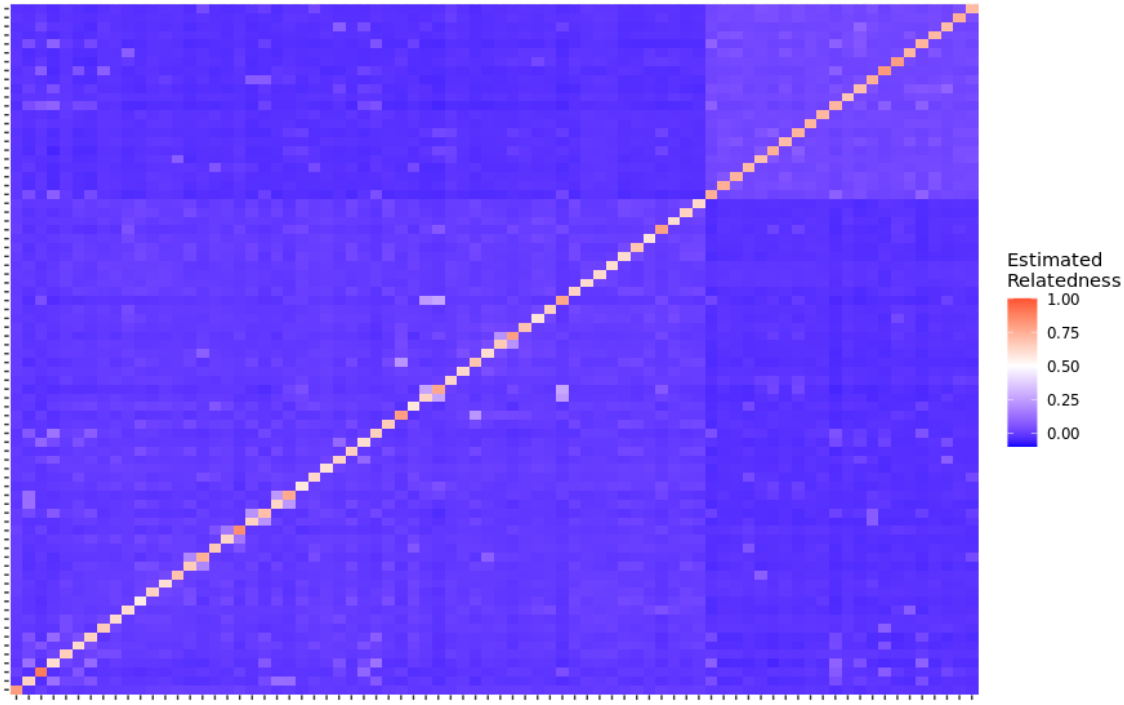
Heat map of estimated relationships across the 78 families using all 214,449 markers VanRaden

### Prediction of Parental Breeding values using a single batch

When predicting with all data, using a covariance matrix based on SNP genotypes had higher predictive accuracy (assessed as the correlation coefficient) than using covariances based transcript abundance or on a combination of SNP genotypes and transcript abundance, using either OmicKriging (P=3.748e-3) or EN (P=1.418e-3). (Table 1) While the correlation was slightly higher in EN, the paired t-test of absolute residual means when using SNPs or transcripts was not statistically significant (P=0.202), suggesting that the two statistical methods are not different in overall accuracy of prediction. We compared prediction accuracies from data prepared using two different thresholds of removing transcripts which had low expression (mean of four counts within family versus a mean of eight counts) and concluded that predictions were similar enough with no residual differences; therefore, remaining analyses were conducted using a threshold of eight counts, or a log-2 transformed family mean count of three (Supplementary Table 2).

**Table 1.** Correlation (r) values of prediction vs. actual breeding values of families in batch two (N=11) and three (N=22) using two software (OmicKriging, EN). RMSE in parentheses and residuals significantly less P<.05 than ALL* or setA+. ^ Indicates linear regression p-value less than 0.05

| Filter Type | OmicKriging |  |  | EN |  |  |
| --- | --- | --- | --- | --- | --- | --- |
|  | *SNPs* | *Txpts* | *Combined* | *SNPs* | *Txpts* | *Combined* |
| ALL | **0.49<sup>^</sup>**<br>**(15.89)** | 0.18<br>(19.27) | 0.36<sup>^</sup> (17.13) | **0.53<sup>^</sup>**<br>**(15.33)** | 0.27<br>(17.14) | 0.43<sup>^</sup> (15.92) |
| setA | 0.49<sup>^</sup><br>(15.86) | 0.51<sup>^*</sup><br>(15.10) | **0.57<sup>^\*</sup>**<br>**(15.36)** | 0.63<sup>^*</sup><br>(14.25) | 0.52<sup>^*</sup><br>(14.86) | **0.65<sup>^\*</sup>**<br>**(14.04)** |
| setB | 0.52<sup>^</sup><br>(15.81) | 0.57<sup>^*+</sup><br>(14.59) | **0.63<sup>^\*+</sup>**<br>**(15.04)** | 0.65<sup>^*</sup><br>(14.08) | 0.64<sup>^*+</sup><br>(13.31) | **0.72<sup>^\*</sup>**<br>**(13.28)** |
| setC | 0.67<sup>^</sup><br>(14.96) | 0.54<sup>^*</sup><br>(14.86) | **0.72<sup>^\*+</sup>**<br>**(14.68)** | 0.70<sup>^*</sup><br>(13.96) | 0.62<sup>^*+</sup><br>(13.62) | **0.74<sup>^\*</sup>**<br>**(13.36)** |

The lfdr adjusted p-values from a one-way ANOVA using batch and SNP levels resulted in the removal of only 3,359 features. Utilizing this set of SNPs for prediction had no impact compared to using all SNPs (results not shown). Application of a relative variance threshold > 1 to the data removed 189,841 SNPs with 21,249 remaining for prediction (setA) (Supplementary Figure 2). Compared to using all SNPs, the resulting set significantly reduced residual error of prediction (P=1.316e-2) and improved accuracy using the EN model (P=8.5079e-5); however, use of this subset of SNPs did not significantly reduce residual mean errors in OmicKriging (P=.439) and had relatively the same correlation which was significant (P=3.759e-3) (Table1).

Applying one-way ANOVA to transcripts and batch with lfdr < 0.05 resulted in retaining 1,747 transcripts out of the original 34,942 (setA). Prediction with this reduced set as compared to the baseline resulted in a significant decrease in residual errors within both OmicKriging (P=.02011) and EN (P=.03266) (Table 1). Similarly, the regression of predicted versus true breeding values were significant in both OmicKriging (P=2.268e-3) and EN (P=1.775e-3) (Table 1). The highest correlation among setA predictions was observed by taking the mean of predictions from transcripts and SNPs, for both OmicKriging (P=8.508e-5) and EN (P=4.058e-5) (Table 1).

Although the combination of transcripts and SNPs with EN produced a higher correlation accuracy, a paired t-test of absolute residual means between EN and OmicKriging predictions was marginally significant (P=0.0509) (Table 1).

Removing correlated features from the setA features resulted in retaining 5,644 out of 21,249 SNPs and 890 out of 1,747 transcripts (setB). Compared to setA predictions, removing correlated features did not significantly improve accuracy of prediction using SNPs (Table 1). However, they did improve transcripts, with the highest prediction accuracy using EN (P=5.385e-4) (Table 1). Similar to setA predictions, combining SNP and transcript EN predictions resulted in the highest correlation across all of setB (P=2.210e-6) (Table 1).

The one-way ANOVA estimates comparing the 5,644 SNPs of setB to volume breeding values did not produce adjusted p-values less than P=0.05; therefore, we filtered on unadjusted p-values, which resulted in a total of 236 SNPs (setC). Selection of transcripts correlated greater than r=0.05 with phenotypes retained 668 transcripts (setC). Similar to the previous sets, combining the two data types performed the best in both OmicKriging (P=1.989e-6) and EN (P=1.0413e-6) (Table 1). Despite these being the highest prediction accuracies observed, a paired t-test between absolute residual errors was not statistically significant when compared to setB in either OmicKriging (P=.358) or EN (P=.8768) (Table 1). Overall, the highest correlation was observed combining the two data types using setC with EN (Figure 3).

**Figure 3.**
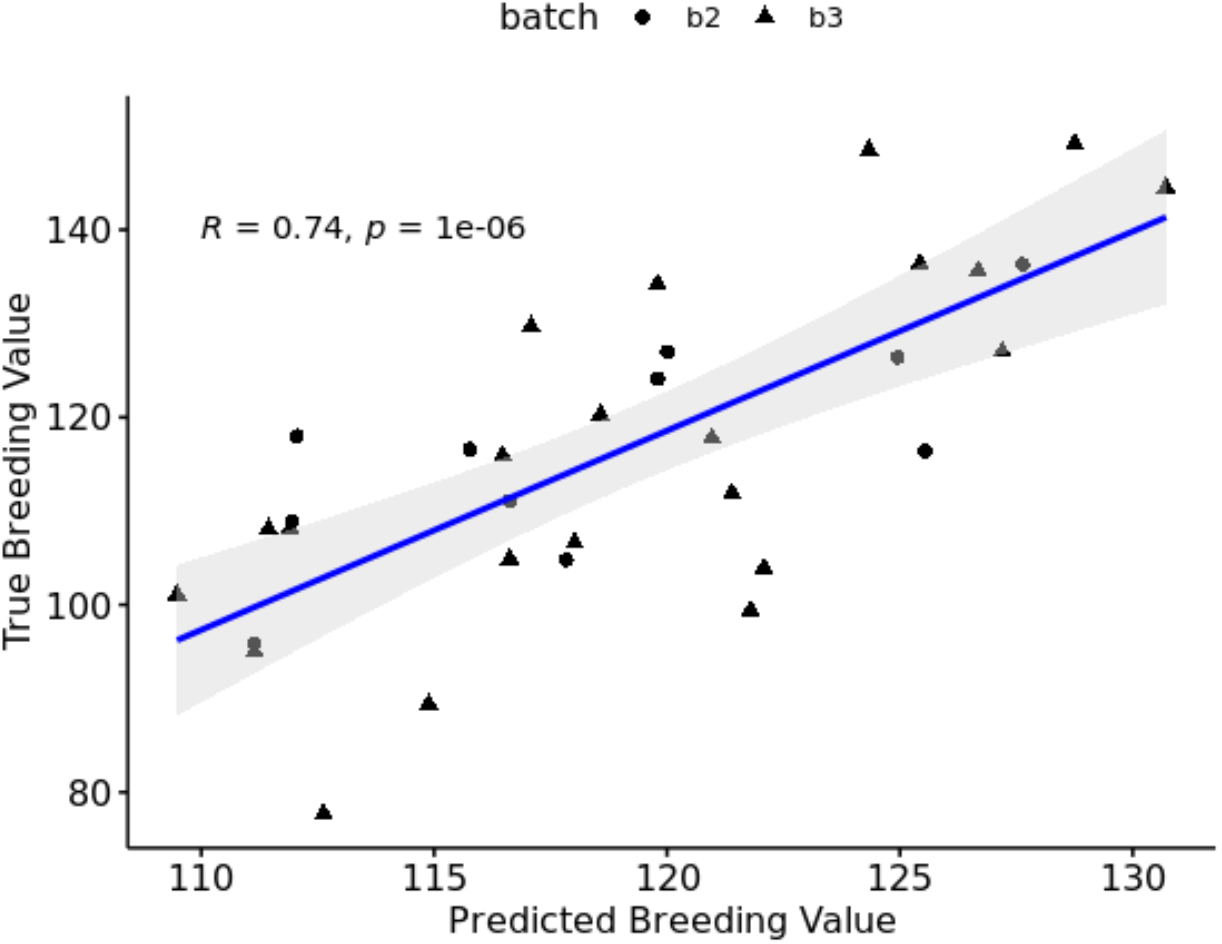
Predicted versus true breeding values of batch two (N=11) and batch three (N=22) families using setC combined predictions with EN.

### Prediction of breeding values using two batches for training

#### Batch Two Predictions

Prediction of batch two (N=11) when training on batch one and three (N=67) resulted in the highest accuracy using all SNPs for both OmicKriging (P=5.079e-3) and EN (P=5.030e-3) (Table 2). While correlations were nearly identical between the two statistical methods, EN had a slightly lower RMSE and higher correlation (Figure 4).

**Table 2.** Correlation (r) values of batch two families (N=11) predicted vs. actual breeding values. RMSE in parentheses. ^ indicates p-value less than 0.05

| Filter Type | OmicKriging |  |  | EN |  |  |
| --- | --- | --- | --- | --- | --- | --- |
|  | *SNPs* | *Txpts* | *Combined* | *SNPs* | *Txpts* | *Combined* |
| ALL | **0.78<sup>^</sup>**<br>**(8.69)** | 0.23<br>(11.35) | 0.54 (9.88) | **0.78<sup>^</sup>**<br>**(8.34)** | 0.60<br>(8.76) | 0.75<sup>^</sup><br>(7.89) |
| setA | **0.64<sup>^</sup>**<br>**(9.16)** | 0.50<br>(9.52) | 0.61<sup>^</sup> (9.06) | **0.62<sup>^</sup>**<br>**(8.76)** | 0.41<br>(10.59) | 0.54<br>(9.22) |
| setB | 0.57<br>(9.53) | 0.47<br>(9.86) | **0.55 (9.20)** | **0.54**<br>**(9.49)** | 0.34<br>(11.64) | 0.46<br>(9.90) |
| setC | 0.08<br>(11.16) | 0.44<br>(10.20) | **0.42 (9.97)** | **0.43**<br>**(10.09)** | 0.36<br>(12.76) | 0.44<br>(10.22) |

**Figure 4.**
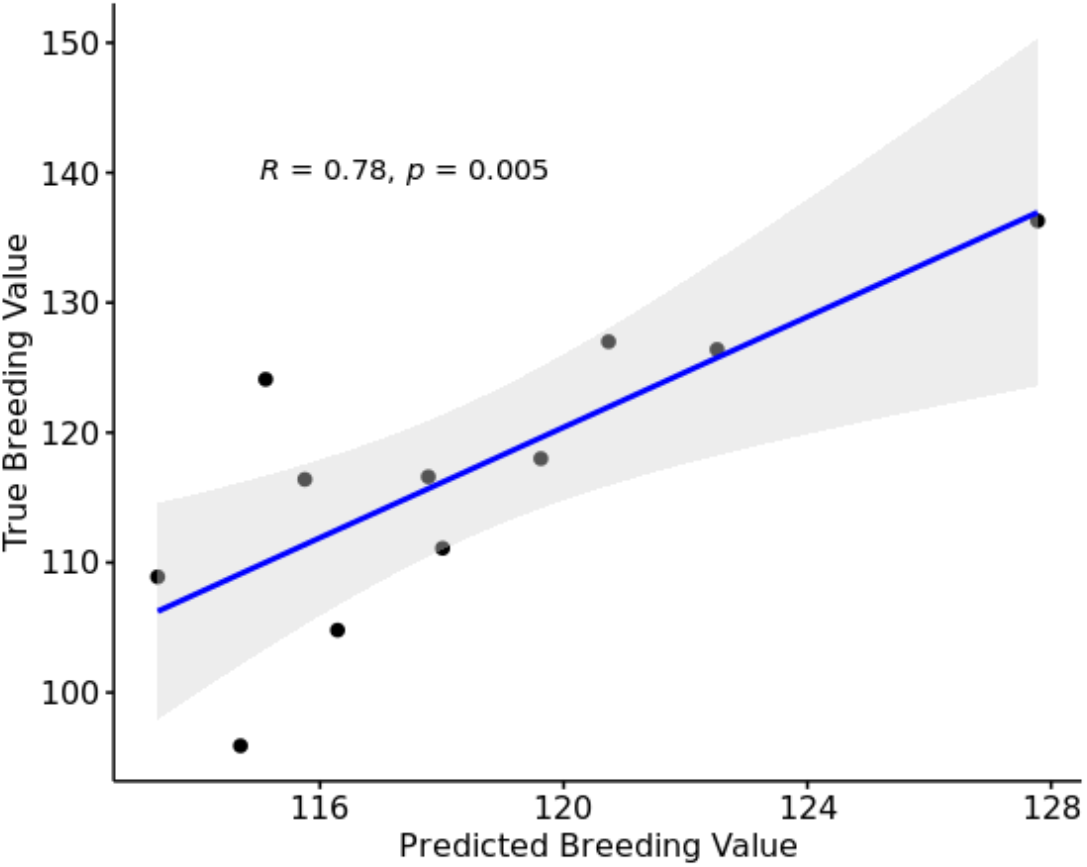
Predicted versus true breeding values of batch two (N=11) families using ALL SNPs with EN.

Utilizing transcripts, prediction accuracies in EN were moderate, however none were significant, ranging from P=0.314482 in setB to P=0.50515 using all data. Similarly, none of the OmicKriging correlations were significant, ranging from P=0.495611 using all data to P=0.115929 using setA (Table 2).

One-way ANOVA estimates comparing the 5,644 SNPs resulted in 333 SNPs, and selection of transcripts correlated greater than r=0.05 with phenotypes retained 613 transcripts (setC). In general, these sets typically performed among the worst within each data type and model (Table 2).

#### Batch 3 Predictions

Across both OmicKriging and EN, the prediction of batch three families (N=22) when training on batch one and two (N=56), generally resulted in the highest accuracies when combining SNPs and transcripts within each filter type (Table 3). In the three cases where using only one data type produced the highest correlation, the combination of transcripts and SNP predictions were nearly equivalent (Table 3).

**Table 3.** Correlation (r) values of prediction vs. actual breeding values of families in batch three (N=22). RMSE in parantheses and residuals significantly less than P<.05 than ALL * or setA+, #setB. ^ Indicates linear regression p-value less than 0.05.

| Filter Type | OmicKriging |  |  | EN |  |  |
| --- | --- | --- | --- | --- | --- | --- |
|  | *SNPs* | *Txpts* | *Combined* | *SNPs* | *Txpts* | *Combined* |
| ALL | 0.44<sup>^</sup><br>(18.13) | 0.43<sup>^</sup><br>(19.84) | **0.45<sup>^</sup>**<br>**(18.68)** | **0.48<sup>^</sup>**<br>(17.50) | 0.40<br>(18.03) | 0.47<sup>^</sup> (17.61) |
| setA | 0.39<br>(18.30) | **0.64<sup>^\*</sup>**<br>**(16.54)** | 0.60<sup>^</sup> (17.34) | 0.48<sup>^</sup><br>(17.10) | 0.55<sup>^*</sup><br>(16.54) | **0.59<sup>^\*</sup>**<br>(16.41) |
| setB | 0.42<br>(18.31) | **0.67<sup>^</sup>**<br>**(16.25)** | 0.66<sup>^</sup> (17.16) | 0.58<sup>^*</sup><br>(16.72) | 0.63<sup>^*</sup><br>(15.52) | **0.71<sup>^\*</sup>**<br>**(15.71)** |
| setC | 0.59<sup>^</sup><br>(17.44) | 0.67<sup>^*</sup><br>(16.88) | **0.79<sup>^\*+ #</sup>**<br>**(16.45)** | 0.64<sup>^*</sup><br>(16.43) | 0.65<sup>^*</sup><br>(15.80) | **0.73<sup>^\*</sup>**<br>**(15.84)** |

Using EN and applying removal of correlated features from SNPs (setB), resulted in significantly improved prediction, significant correlation (P=5.110e-3), and reduced residual errors compared to using all SNP data (P=0.04947) (Table3). Similar to what was observed when predicting both batches two and three, setA predictions with transcripts increased accuracy in both statistical methods and applying the removal of correlated features (setB) further improved predictions when compared to the baseline. (EN: P=7.072e-4 and OK: P=6.559e-4). The highest correlation within setB was observed using EN with the combination of transcripts and SNPs (P=2.351e-4) (Table3).

ANOVA estimates with the 5,644 SNPs resulted in 321 SNPs, and selection of transcripts correlated greater than r=0.05 with phenotypes retained 631 transcripts. Similar to the previous filters, combining the two data types performed the best in OmicKriging (P=1.42791e-5) and EN (P=1.31267e-4) (Table 3). Although the correlation was higher in OmicKriging than EN, the RMSE was also slightly higher but not significantly greater (P=0.1899) (Figure 5).

**Figure 5.**
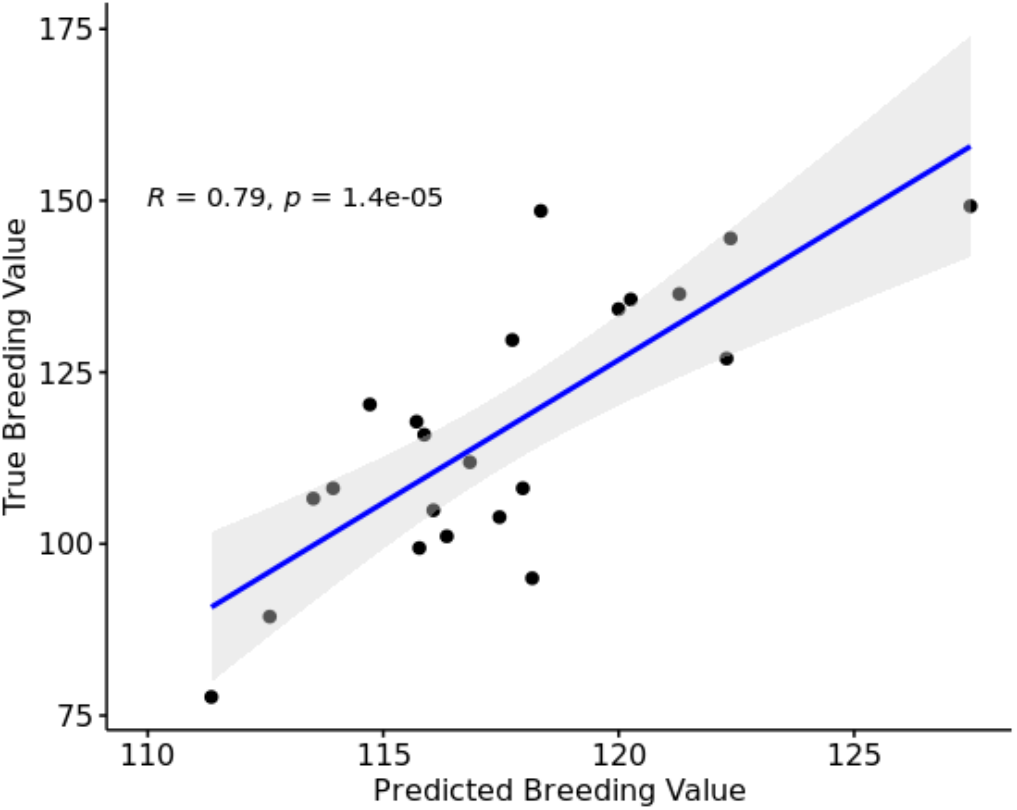
Predicted versus true breeding values of batch three (N=22) families using using setC combined predictions with OmicKriging.

## Discussion

Genomic selection can potentially impact loblolly pine by reducing the length of a given breeding cycle and increasing selection intensity, or genetic gain, made from a cycle [45]. In order to meet that potential, a few hurdles that must be overcome include: its large genome size, significant heterogeneity, and low amount of linkage disequilibrium [46]. Given these difficulties, it may be a while before GS can provide the same benefits to forest tree breeding as it has to annual crops or livestock species with more population structure, deeper pedigrees, and less overall genetic diversity. While waiting for loblolly pine genomic selection, exploring other opportunities to leverage ‘omic’ technologies should be of interest.

In this study, we proposed the idea of enabling transcriptomic selection through the collection of RNA-sequencing data, and tested the concept utilizing three batches of seedlings produced from loblolly pine families spanning a range of phenotypic values for stem volume productivity. RNA-sequencing allows the collection of gene expression patterns and sequence variation within a single experiment; while arrays can only provide either gene expression levels or SNP genotypes, they cannot provide both [47]. One novel aspect of transcriptomic selection carried out in this experiment was aligning reads to a transcriptome reference rather than the genome reference and averaging the genotypes across biological replicates. In theory, analyzing the transcriptomic data in this way allows us to average across members of multi-gene families while potentially removing some environmental differences in gene expression.

When evaluating gene expression reproducibility, biological replicate transcript information showed a high degree of similarity. These results agree with previous studies showing the high reproducibility of expression estimates [48,49]. There was also a high similarity among replicates across batches, which provides confidence in reproducing certain expression levels across years and experiments if this method were used in the future. While family-mean expression levels were similar among families, the SNP relationship matrix showed minimal relationships among the families sequenced. This observation of high similarity of gene expression levels among families suggests that some genes may be expressed in the seedling stage as part of development independent of any family influence [50].

The lack of relationships in the SNP matrix makes this study unusual compared to most other genomic prediction experiments. Most previous omic prediction studies within row crops have relied on pre-existing relationships among the individuals sequenced [51]. Additionally, most forest tree genomic predictions have been evaluated using relatively close relationships between training and test populations. [52,53]. Provided the evidence that both gene expression and phenotype are heritable, it is reasonable to propose transcriptomics has some value in understanding differences in phenotypes among individuals [54,55].

When training with a single batch, both the OmicKriging and EN models displayed relatively similar accuracies when using a single data type (SNPs or transcripts) in combination with a specific filter type. This observation tends to agree with previous genomic selection studies assessing ridge regression and elastic net [56, 57], which concluded that the shrinkage of estimates does not significantly impact prediction accuracy when evaluating complex polygenic traits. More important than the model was the inclusion of certain variables. Prediction with variables that showed variation among families and were not highly correlated with one another seemed to have the largest impact compared to using all data. This observation makes sense given that EN has the potential to either include or remove a set of correlated variables [58]. Collinearity of the variables used in the correlation matrix could skew assumed relationships among individuals [59]. Surprisingly, in both OmicKriging and EN, the combination of SNPs and transcripts outperformed using either alone, suggesting that there may be different types of information from the two contributing to the prediction accuracy. Similar findings have been reported in maize, where prediction of dry matter yield and all other traits evaluated saw a slight boost in prediction accuracies when combining genotype and transcript information to predict hybrid phenotype [51].

In theory, combining batches should provide higher prediction accuracies on the left-out test population used for prediction. Training using batch one and two to predict batch three gave a similar result as when predicting using a model trained only with batch one. However, the prediction of batch two families when using batch one and three as training gave a dramatically different outcome. Using all the data, without filtering for variation among families or association with phenotype, came close to the types of other accuracies seen, and SNPs seemed to outperform transcript levels or both SNPs and transcripts together. The reason for the observed poor prediction accuracy of batch two using models trained on batch one and three is unknown. However, this observation may be due to sampling effects of families, because batch two contains only 11 families not present in batch one or three. It may be that by chance those 11 families are different in some way from the families in batch three, so that models trained with batch one alone predict batch two much more accurately than do models trained on batches one and three. A recent assessment of prediction ability within a population of clonally replicated loblolly pine saw close to a 50% reduction in prediction ability if removing full-sib relatives from the training population [60]. This observation also may be because batch one and two were sequenced with different instruments, using libraries prepared by different centers, or sequencing was done with different providers. The SNP relationship matrix shows a discernable differentiation between batch one/two and batch three, suggesting that these technical differences had some effect on the detection and genotyping of SNP loci. Further experiments will be required to determine the relative contribution of any of the above factors.

Genomic selection is generally applied as a form of forward selection. Every offspring is individually-genotyped, and then those individual genotypes are analyzed, with or without phenotypes, to decide which individual to select for further breeding. Our transcriptomic selection approach is different; instead of genotyping individual offspring separately, we sequence them as a pool. The unit of selection is not a seedling from the genotyped set but the parent of the genotyped seedlings. Despite the partial inconsistency in how a filter type performed in prediction, overall accuracies suggest this method may potentially be included as a form of very early backward selection, as it applies to the current loblolly pine breeding approach. In principle, both methods could be used together in the same breeding program by first using individual genotyping and genomic prediction to make forward selections. Those forward selections could then be top-grafted and allowed to open-pollinate the first crop of female strobili produced. Those OP cones would be collected as soon as three years after top-grafting, and RNA-seq experiments conducted on a sample of the seedlings to perform backward selection among the forward selections. The remaining OP seedlings could also be planted in field trials to get additional phenotypic data to update the models used to train the genomic and transcriptomic selection models.

Further improvement could be made in the application of transcriptomic prediction by leveraging megagametophytes instead of the seedlings themselves. The megagametophyte is the nutritive tissue of the conifer seed, analogous to the endosperm of angiosperm seeds, but unlike the endosperm, the megagametophyte is derived from the same haploid product of maternal meiosis as the embryo. Preliminary data from a single family suggests that RNA-seq with RNA from megagametophytes yields results very similar to those obtained with RNA from seedlings, and may work just as well as RNA from growing seedlings. The ability to utilize megagametophytes would hasten analysis by saving the time required for seed stratification and seedling growth. Additionally, it may reduce the potential for batch effects related to greenhouse growth conditions. If using seedlings in future experiments, it may be important to either utilize a balanced design or have a control family grown across batches to ensure that sequencing differences, whether due to growth conditions or to the methods or sequencing technology used, are not severely inflating the batch effect.

Overall, the findings from this study have the potential to provide a significant impact on how loblolly pine breeding populations are managed. Transcriptomic selection within loblolly pine would allow for future selections to be screened earlier, and allow greater selection intensity on breeding populations while also potentially providing more gain to be made over time. The exploration of using other omic technologies, besides strictly genomic selection, should continue to be of interest to loblolly pine breeders and potentially used in conjunction with genomic selection to provide the best crop moving forward.

