## Supplemental Material for "Transcriptomic Prediction of Breeding Values in Loblolly Pine"

### Supplementary Material

**Supplementary Table 1.** Family identifier with the corresponding number of biological replicates in each respective batch and type of pollen mix.

| Family Identifier | # Biological Replicates | Batch | OP/PMX/FS |
|---|---|---|---|
| TIP1112568 | 1 | 1 | OP |
| TIP119519 | 1 | 1 | OP |
| TIP143184 | 1 | 1 | OP |
| TIP2291953 x TIP11257736 | 1 | 1 | FS |
| TIP2131691 x TIP2106476 | 2 | 1 | FS |
| TIP1420947 | 2 | 1 | OP |
| TIP1212177 | 2 | 1 | PMX |
| TIP1103086 | 2 | 1 | PMX |
| TIP2194953 | 2 | 1 | OP |
| TIP1119498 | 3 | 1 | OP |
| TIP11031586 | 3 | 1 | OP |
| TIP1123708 x UC | 3 | 1 | PMX |
| TIP1121708 | 3 | 1 | OP |
| TIP11255033 | 3 | 1 | OP |
| IP1212177 x TIP1412064 | 3 | 1 | FS |
| TIP1164622 | 3 | 1 | PMX |
| TIP1190138 | 3 | 1 | PMX |
| TIP1506888 | 3 | 1 | OP |
| TIP1940090 | 3 | 1 | OP |
| TIP21206990 | 3 | 1 | PMX |
| TIP2131691 | 3 | 1 | OP |
| TIP21431566 | 3 | 1 | PMX |
| TIP2211991 | 3 | 1 | OP |
| TIP2291953 | 3 | 1 | OP |
| TIP1420947 x B48 | 3 | 1 | FS |
| TIP1212177 x TIP1412064 | 3 | 1 | FS |
| TIP2387750 | 3 | 1 | OP |
| TIP2446494 | 3 | 1 | OP |
| TIP2448804 | 3 | 1 | OP |
| TIP2523434 | 3 | 1 | PMX |

**Table 1** (continued)

| | | | |
|---|---|---|---|
| TIP2791624 | 3 | 1 | PMX |
| TIP292525 | 3 | 1 | PMX |
| TIP1123708 | 4 | 1 | OP |
| TIP11319300 | 4 | 1 | PMX |
| TIP1142759 | 4 | 1 | OP |
| TIP1242079 | 4 | 1 | OP |
| TIP1281198 | 4 | 1 | OP |
| TIP138245 | 4 | 1 | OP |
| TIP1428572xBA2L20 | 4 | 1 | FS |
| TIP192193 | 4 | 1 | OP |
| TIP2232855 | 4 | 1 | PMX |
| TIP139150 | 5 | 1 | PMX |
| TIP2232855 x C21B | 5 | 1 | FS |
| TIP1306258 | 5 | 1 | OP |
| TIP1428572 | 5 | 1 | PMX |
| TIP1633160 | 5 | 1 | OP |
| WG7478 | 1 | 2 | OP |
| TIP138245 | 1 | 2 | OP |
| TIP143184 | 1 | 2 | OP |
| TIP181745 | 2 | 2 | OP |
| TIP1158330 | 3 | 2 | OP |
| TIP1170385 | 3 | 2 | OP |
| TIP1857822 | 3 | 2 | OP |
| TIP1932699 | 3 | 2 | OP |
| TIP1973363 | 3 | 2 | OP |
| TIP21206990 | 3 | 2 | OP |
| TIP2523434 | 3 | 2 | OP |
| TIP2751739 | 3 | 2 | OP |
| TIP2818013 | 3 | 2 | OP |
| LSG111 | 3 | 2 | OP |
| WG1809 | 3 | 2 | OP |
| WG4302 | 3 | 2 | OP |
| WG4501 | 3 | 2 | OP |
| WG4801 | 3 | 2 | OP |
| WG7192 | 3 | 2 | OP |
| WG7206 | 3 | 2 | OP |
| WG7442 | 3 | 2 | OP |

**Table 1** (continued)

| | | | |
|---|---|---|---|
| WG7720 | 3 | 2 | OP |
| TIP139150 | 4 | 2 | OP |
| TIP163066 | 4 | 2 | OP |
| TIP216059 | 4 | 2 | OP |
| WG7083 | 4 | 2 | OP |
| TIP1685530 | 5 | 2 | OP |
| TIP11065116 | 3 | 3 | OP |
| TIP1145480 | 3 | 3 | OP |
| TIP1253401 | 3 | 3 | OP |
| TIP1256535 | 3 | 3 | PMX |
| TIP150187 | 3 | 3 | PMX |
| TIP1711512 | 3 | 3 | PMX |
| TIP178908 | 3 | 3 | PMX |
| TIP21066782 | 3 | 3 | OP |
| TIP2109755 | 3 | 3 | PMX |
| TIP2120526 | 3 | 3 | PMX |
| TIP21687888 | 3 | 3 | OP |
| TIP2170002 | 3 | 3 | OP |
| TIP2178058 | 3 | 3 | OP |
| TIP224675 | 3 | 3 | OP |
| TIP2277451 | 3 | 3 | OP |
| TIP2306224 | 3 | 3 | OP |
| TIP2539112 | 3 | 3 | OP |
| TIP261659 | 3 | 3 | OP |
| TIP275099 | 3 | 3 | OP |
| TIP2867690 | 3 | 3 | OP |
| TIP3208051 | 3 | 3 | OP |
| TIP330244 | 3 | 3 | OP |

**Supplementary Table 2.** Correlation (r) values of prediction vs. actual breeding values of families in batch two and three using log(4) threshold. RMSE in parentheses and residuals significantly less P<.05 than ALL* or setA+, ^ Indicates linear regression p-value less than 0.05

| | OmicKriging | | | EN | | |
|---|---|---|---|---|---|---|
| Filter Type | *SNPs* | *Txpts* | *Combined* | *SNPs* | *Txpts* | *Combined* |
| ALL | **0.491^** **(15.89)** | 0.158 (18.21) | 0.339^ (16.75) | **0.533^** **(15.33)** | 0.076 (17.87) | 0.324^ (16.23) |
| setA | 0.490^ (15.86) | 0.474^* (15.53) | **0.570^** **(15.59)** | 0.630^* (14.25) | 0.333^* (16.06) | 0.588^* (14.75) |
| setB | 0.518^ (15.81) | 0.639^*+ (14.90) | **0.704^*+** **(15.25)** | 0.653^* (14.08) | 0.528^*+ (14.49) | 0.710^* (13.83) |
| setC | 0.666^ (14.96) | 0.637^*+ (14.97) | **0.776^*+** **(14.80)** | 0.697^* (13.96) | 0.521^*+ (14.60) | 0.713^* (13.85) |

**(A)**

| ***LANE*** | 1 | 2 | 3 | 4 | 5 | 6 | 7 | 8 | 9 | 10 | 11 | 12 | 13 | 14 | 15 | [illegible] |
|---|---|---|---|---|---|---|---|---|---|---|---|---|---|---|---|---|
| ***REP*** | X1 | X2 | X3 | X4 | X5 | X6 | X1 | X2 | X3 | X4 | X5 | X6 | X1 | X2 | X3 | [illegible] |
| **1** | 17a | 133a | 104a | 56a | 31a | 95a | 95a | 17a | 133a | 104a | 56a | 31a | 31a | 95a | 56a | [illegible] |
| **2** | 72b | 6b | 101b | 32b | 132b | 94b | 94b | 132b | 6b | 101b | 32b | 72b | 101b | 32b | 72b | [illegible] |
| **3** | 136c | 67c | 78c | 113c | 46c | 7c | 7c | 78c | 46c | 67c | 136c | 113c | 46c | 67c | 136c | [illegible] |
| **4** | 37d | 123d | 63d | 112d | 91d | 18d | 18d | 123d | 91d | 37d | 112d | 63d | 18d | 123d | 37d | [illegible] |
| **5** | 129e | 106e | 57e | 77e | 21e | 25e | 129e | 77e | 21e | 106e | 25e | 57e | 25e | 106e | 21e | [illegible] |
| **6** | 68f | 13f | 93f | 130f | 38f | 107f | 107f | 68f | 38f | 13f | 130f | 93f | 93f | 13f | 68f | [illegible] |
| **7** | 19g | 34g | 62g | 86g | 108g | 142g | 62g | 19g | 108g | 86g | 34g | 142g | 86g | 19g | 34g | [illegible] |
| **8** | 127h | 75h | 110h | 59h | 20h | 28h | 75h | 59h | 127h | 110h | 28h | 20h | 59h | 28h | 127h | [illegible] |
| **9** | 120i | 61i | 134i | 44i | 14i | 88i | 61i | 120i | 14i | 44i | 134i | 88i | 120i | 14i | 44i | [illegible] |
| **10** | 33j | 4j | 70j | 137j | 92j | 103j | 4j | 70j | 92j | 33j | 103j | 137j | 137j | 70j | 33j | [illegible] |
| **11** | 102k | 79k | 66k | 9k | 139k | 35k | 139k | 35k | 9k | 66k | 79k | 102k | 79k | 102k | 9k | [illegible] |
| **12** | 105l | 125l | 8l | 96l | 48l | 51l | 125l | 48l | 8l | 105l | 96l | 51l | 48l | 96l | 125l | [illegible] |
| **13** | 36m | 60m | 98m | 87m | 131m | 15m | 60m | 87m | 15m | 36m | 98m | 131m | 60m | 15m | 36m | [illegible] |
| **14** | 23n | 65n | 40n | 76n | 100n | 122n | 122n | 23n | 40n | 65n | 76n | 100n | 100n | 76n | 122n | [illegible] |
| **15** | 143o | 119o | 89o | 1o | 45o | 58o | 143o | 89o | 58o | 45o | 119o | 1o | 143o | 58o | 89o | [illegible] |
| **16** | 10p | 64p | 80p | 27p | 97p | 124p | 80p | 64p | 124p | 10p | 97p | 27p | 64p | 97p | 80p | [illegible] |
| **17** | 47q | 2q | 141q | 118q | 55q | 90q | 118q | 90q | 47q | 2q | 55q | 141q | 2q | 141q | 55q | [illegible] |
| **18** | 114r | 43r | 54r | 84r | 126r | 22r | 126r | 54r | 22r | 43r | 114r | 84r | 84r | 22r | 54r | [illegible] |
| **19** | 111s | 3s | 30s | 83s | 138s | 49s | 138s | 111s | 30s | 49s | 83s | 3s | 49s | 30s | 3s | [illegible] |
| **20** | 24t | 135t | 82t | 99t | 39t | 50t | 82t | 39t | 24t | 135t | 99t | 50t | 99t | 82t | 24t | [illegible] |
| **21** | 109u | 140u | 69u | 85u | 26u | 12u | 85u | 26u | 69u | 109u | 140u | 12u | 12u | 69u | 85u | [illegible] |
| **22** | 41v | 117v | 5v | 53v | 128v | 74v | 117v | 128v | 53v | 74v | 5v | 41v | 5v | 53v | 128v | [illegible] |
| **23** | 115w | 16w | 42w | 144w | 52w | 81w | 42w | 81w | 115w | 52w | 144w | 16w | 52w | 115w | 42w | [illegible] |
| **24** | 116x | 29x | 71x | 121x | 11x | 73x | 73x | 116x | 71x | 29x | 11x | 121x | 116x | 11x | 29x | [illegible] |

**Supplementary Figure 1.** Experimental design of sequencing layout for all technical replicate RNA samples from (A) Batch 1 and (B) Batch 2.

RESTRICTED

**Supplementary Figure 1** (continued)

**(B)**

| *LANE* | 1 | 2 | 3 | 4 | 5 | 6 | 7 | 8 | 9 | 10 | 11 | 12 |
|---|---|---|---|---|---|---|---|---|---|---|---|---|
| ***REP*** | X1 | X2 | X3 | X4 | X5 | X6 | X1 | X2 | X3 | X4 | X5 | X6 |
| **1** | 32a | 10a | 86a | 49a | 49a | 86a | 32a | 10a | 32a | 10a | 86a | 49a |
| **2** | 15b | 60b | 25b | 92b | 15b | 92b | 25b | 60b | 25b | 92b | 15b | 60b |
| **3** | 57c | 3c | 28c | 85c | 28c | 85c | 3c | 57c | 28c | 85c | 57c | 3c |
| **4** | 66d | 11d | 91d | 31d | 31d | 66d | 91d | 11d | 91d | 31d | 66d | 11d |
| **5** | 24e | 84e | 43e | 69e | 69e | 24e | 43e | 84e | 43e | 84e | 24e | 69e |
| **6** | 64f | 93f | 13f | 34f | 13f | 93f | 34f | 64f | 93f | 34f | 64f | 13f |
| **7** | 53g | 22g | 74g | 42g | 42g | 74g | 22g | 53g | 74g | 42g | 53g | 22g |
| **8** | 7h | 68h | 26h | 83h | 7h | 26h | 83h | 68h | 26h | 68h | 7h | 83h |
| **9** | 16i | 45i | 75i | 58i | 16i | 45i | 75i | 58i | 16i | 45i | 58i | 75i |
| **10** | 37j | 59j | 80j | 1j | 1j | 37j | 59j | 80j | 80j | 1j | 59j | 37j |
| **11** | 48k | 95k | 8k | 54k | 54k | 8k | 95k | 48k | 95k | 48k | 54k | 8k |
| **12** | 96l | 36l | 12l | 50l | 36l | 12l | 96l | 50l | 36l | 50l | 12l | 96l |
| **13** | 78m | 18m | 61m | 46m | 61m | 46m | 78m | 18m | 61m | 46m | 18m | 78m |
| **14** | 56n | 88n | 4n | 47n | 47n | 56n | 4n | 88n | 56n | 4n | 47n | 88n |
| **15** | 90o | 5o | 65o | 38o | 38o | 5o | 90o | 65o | 5o | 90o | 38o | 65o |
| **16** | 6p | 77p | 33p | 67p | 77p | 67p | 6p | 33p | 6p | 33p | 67p | 77p |
| **17** | 89q | 35q | 55q | 21q | 89q | 55q | 35q | 21q | 89q | 21q | 35q | 55q |
| **18** | 81r | 29r | 52r | 2r | 52r | 29r | 81r | 2r | 2r | 81r | 29r | 52r |
| **19** | 17s | 40s | 79s | 72s | 72s | 17s | 40s | 79s | 79s | 17s | 72s | 40s |
| **20** | 76t | 62t | 19t | 27t | 19t | 27t | 62t | 76t | 19t | 62t | 27t | 76t |
| **21** | 71u | 94u | 30u | 14u | 14u | 71u | 94u | 30u | 14u | 30u | 94u | 71u |
| **22** | 44v | 63v | 87v | 20v | 63v | 44v | 87v | 20v | 87v | 20v | 44v | 63v |
| **23** | 41w | 51w | 82w | 23w | 82w | 41w | 23w | 51w | 41w | 51w | 82w | 23w |
| **24** | 73x | 9x | 70x | 39x | 9x | 73x | 39x | 70x | 73x | 9x | 39x | 70x |

RESTRICTED

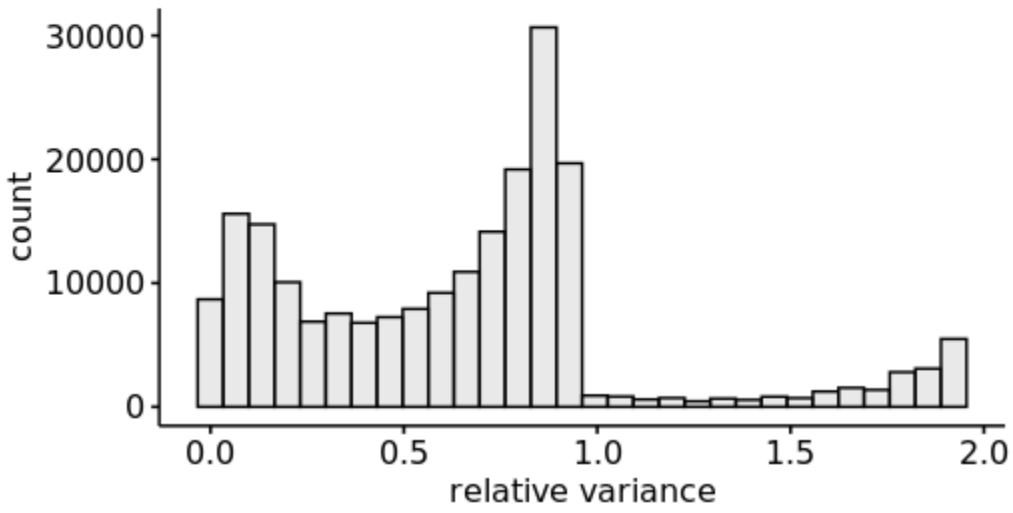


**Supplementary Figure 2.** Histogram of relative variance (variance/mean) values for each of the 211,090 SNP loci across all 78 families.

RESTRICTED
